# scYeast: a Biological-knowledge-guided Foundation Model on Yeast Single-Cell Transcriptomics

**DOI:** 10.1101/2025.08.20.671179

**Authors:** Xingcun Fan, Wenbin Liao, Luchi Xiao, Xuefeng Yan, Hongzhong Lu

## Abstract

Though large-scale pre-trained models are vital for foundational cell modeling, most focus on human or mouse systems, neglecting model organisms like yeast (*Saccharomyces cerevisiae*) and often failing to use existing biological prior knowledge effectively. To address this, we present scYeast, the first foundational cell model tailored for yeast that seamlessly embeds biological priors. scYeast employs a novel asymmetric parallel architecture to infuse transcriptional regulatory information directly into the Transformer’s attention mechanism, leveraging established biological knowledge during training. Pre-trained on large-scale yeast single-cell transcriptomics data, scYeast demonstrates strong generalization and biological interpretability. It excels in zero-shot tasks, such as inferring regulatory relationships and identifying critical cell states. After fine-tuning, scYeast performs exceptionally across a diverse set of tasks from cell type classification to predicting growth doubling time and gene perturbation response. Additionally, using transfer learning, scYeast can be adapted to other omics datasets, such as proteomics, thus broadening its utility in big data analysis. Overall, scYeast is a powerful tool for yeast single-cell biology research and sets a new standard for integrating foundational models with biological prior knowledge, dramatically accelerating the pace of discovery in yeast synthetic and systems biology and providing a replicable framework for other organisms.

## Introduction

Recently, foundation models, typically built on the Transformer architecture at their core, have revolutionized the research in natural language processing and life sciences ^1^. By leveraging self-supervised learning frameworks and large-scale parameterization, these models can extract abstract, reusable representations from enormous, intricate datasets. Crucially, these models can be employed to capture and simulate the whole cellular metabolic activities, thus making it possible to build virtual cell models at unprecedented scales ^2,3^.

The widespread application of single-cell sequencing technologies has led to the accumulation of massive volumes of single-cell transcriptomics data, providing chances for the reconstruction of large-scale foundation models. For instance, scBERT ^4^ developed a pre-trained deep neural network model, adopting the pre-training and fine-tuning methodology of BERT. It gains a general understanding of gene-gene interactions through pre-training on a vast amount of unlabeled scRNA-seq data, which is then transferred and fine-tuned for supervised cell type annotation tasks on unseen scRNA-seq data. scFoundation ^5^ developed an asymmetric Transformer-style architecture and pre-training task design, encompassing approximately 20,000 genes and pre-trained on over 50 million human single-cell transcriptome profiles, enabling it to effectively capture the complex contextual relationships between genes across various cell types and states. DeepIMAGER ^6^ infers cell-specific regulatory networks through deep learning and data integration, employing a supervised learning approach that converts the co-expression patterns of gene pairs into image-like representations and utilizes transcription factor (TF) binding information for model training. Geneformer ^7^ introduced a modified Transformer encoder with a latent array to eliminate dependency on nucleotide sequences. DeepRIG ^8^ proposed a graph-based deep learning model that constructs a prior regulatory graph by transforming the gene expression patterns of the data into co-expression patterns. It then utilizes a graph autoencoder model to embed the global regulatory information contained in the graph into latent gene embeddings, thereby reconstructing the gene regulatory network.

However, despite the significant advancements that deep learning has brought to high-dimensional modeling and the discovery of regulatory mechanisms from single-cell data, existing methods still fall short in the deep integration of biological prior knowledge. The majority of current studies merely embed prior knowledge as a form of multi-modal data, concatenating it with input features to be fed into the model, which fails to directly demonstrate the effective utilization of this knowledge during the model’s inference process. For example, GRNPT, a novel Transformer-based framework, leverages textual descriptions of individual genes from the NCBI database, generating embedding vectors via a large language model to capture biological information, which are then used as part of the input data for inferring regulatory relationships ^9^. GeneCompass integrates four types of prior biological knowledge—gene connectivity networks, promoter information, gene family annotations, and co-expression relationships—by creating separate embeddings for each category and combining them into a joint input vector for training a deep learning model ^10^. Similarly, DNA sequences, protein sequences, and gene descriptions have also been separately embedded to predict intergenic relationships ^11^. It is evident from these approaches that prior knowledge is treated as a multi-modal input, where different forms of information are independently embedded, concatenated into a new vector, and then used for model training and prediction. Analogous methods for integrating multi-modal data are also employed in other research areas ^12-14^. Consequently, when faced with complex biological scenarios, the interpretability of existing large models and the efficacy of their biological knowledge utilization remain limited.

As a crucial model organism, *Saccharomyces cerevisiae* is extensively used in fields of systems and synthetic biology ^15-17^. Meanwhile, a wealth of prior knowledge and omics datasets have been accumulated for *S. cerevisiae* ^18-22^. However, most current foundation models are built for human or murine systems. Therefore, developing a foundational model specially for yeast holds significant scientific value.

Here, we propose scYeast, a foundational model for yeast based on the deep fusion of prior knowledge and large-scale single-cell/bulk transcriptomic datasets. Inspired by DSGraphformer ^23^, scYeast introduces an asymmetric parallel transformer architecture that deeply integrates transcriptional regulatory prior knowledge with the attention computation mechanism. This design ensures that scYeast thoroughly considers the incorporated prior knowledge throughout its training process. The model is first pre-trained in a self-supervised manner on a large corpus of unlabeled single-cell transcriptomics data. Subsequently, it can be adapted to a diverse array of downstream tasks, i.e., the inference of condition-specific transcriptional regulatory relationships and classification of cell states. Furthermore, the transfer learning was employed to enlarge the applications of scYeast with additional proteomic data as input. scYeast not only provides a novel tool for investigating the functional states and phenotypic variations of yeast cells but also presents a new research paradigm for the deep integration of foundational models with biological knowledge.

## Results

### Framework of scYeast

scYeast is a transformer-based pre-trained large model designed to construct a foundational model of the yeast Transcriptional Regulatory Network (TRN) that deeply integrates prior knowledge. Drawing inspiration from the static graph concept of DSGraphformer ^23^, scYeast employs an asymmetric parallel deep learning architecture. This design allows for the introduction of prior transcriptional regulatory relationships, in the form of an adjacency matrix, directly into the attention computation process. Consequently, the model can fully leverage this prior knowledge during computation and training. The overall architecture of scYeast is depicted in Fig. 1a. The model begins by embedding single-cell transcriptomics data into high-dimensional vectors through expression token embedding. Concurrently, gene co-expression relationships are inferred from high-quality pan-transcriptome data ^24^, using a Spearman correlation coefficient greater than 0.5 as the threshold. Based on these inferred co-expression relationships, a name embedding vector is generated for each gene after an embedding step. Subsequently, the expression token embeddings are randomly masked and then summed with the gene name embeddings before being fed into two parallel encoder modules.

**Figure 1.**
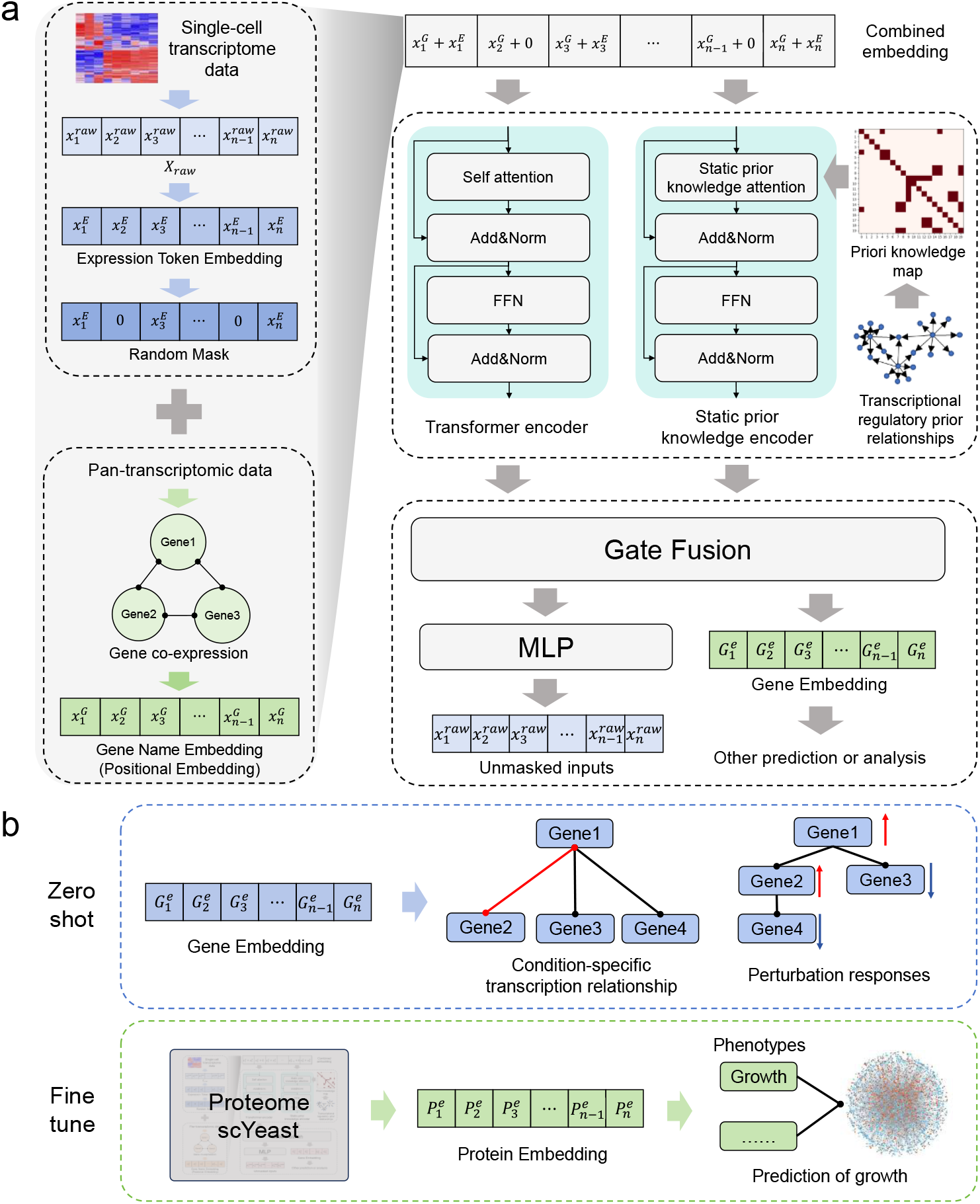
Framework and conceptual diagram of scYeast. (a) The overall framework of scYeast. It employs a parallel architecture to introduce prior knowledge, in the form of an adjacency matrix, into the attention mechanism, achieving a deep fusion of mechanistic knowledge. (b) scYeast can perform various prediction and analysis tasks, such as cell state classification and gene perturbation prediction, through either fixed-parameter or fine-tuning approaches. It can also be transferred to proteomics data to enable additional functions, including cell growth prediction.

Within scYeast (Fig. 1a), two parallel Transformer encoders collaboratively undertake the task for feature extraction, yet they focus on distinct sources of signals. The first branch employs the standard multi-head Query-Key-Value (QKV) attention mechanism, consistent with models like BERT. This “dynamic” encoder relies entirely on the single-cell transcriptomics data itself to capture potential novel gene expression dependencies that may emerge under varying conditions. In parallel, the “static” encoder explicitly injects mechanistic prior knowledge (a transcription factor–target gene adjacency matrix) at the attention layer. After both encoders complete their respective contextual encoding, their outputs are passed to a learnable gating fusion unit. The fused representation is then processed through a Multilayer Perceptron (MLP) to reconstruct the original, unmasked input vector, thereby accomplishing the self-supervised pre-training (Methods).

Leveraging the parameter representations extracted from the pre-trained model and subsequent fine-tuning, scYeast can be broadly applied to yeast large-scale data analysis, including integrative analyses linking omics data with phenotypic traits (Fig. 1b). In scYeast, after pre-training, all model parameters can be frozen; when new transcriptomic data are input, the output of the gate fusion module can be interpreted as a gene embedding that effectively integrates both data-driven features and prior biological knowledge. These embeddings can then be readily employed for diverse downstream analytical or predictive tasks (Fig. 1b).

### scYeast can analyze condition-specific transcriptional regulatory relationships

The scYeast could effectively leverage prior knowledge combined with information learned from massive pre-training datasets to generate high-dimensional gene embeddings that encapsulate both gene expression levels and regulatory information. This lays a solid foundation for in-depth investigation of transcriptional regulatory relationships. Using single-cell transcriptomics data from the yeast cells under different aging condition (young, early-aged, and late-aged) ^25^, we employed scYeast to probe the dynamic changes in intracellular gene regulatory relationships across different age stages (Fig. 2a).

**Figure 2.**
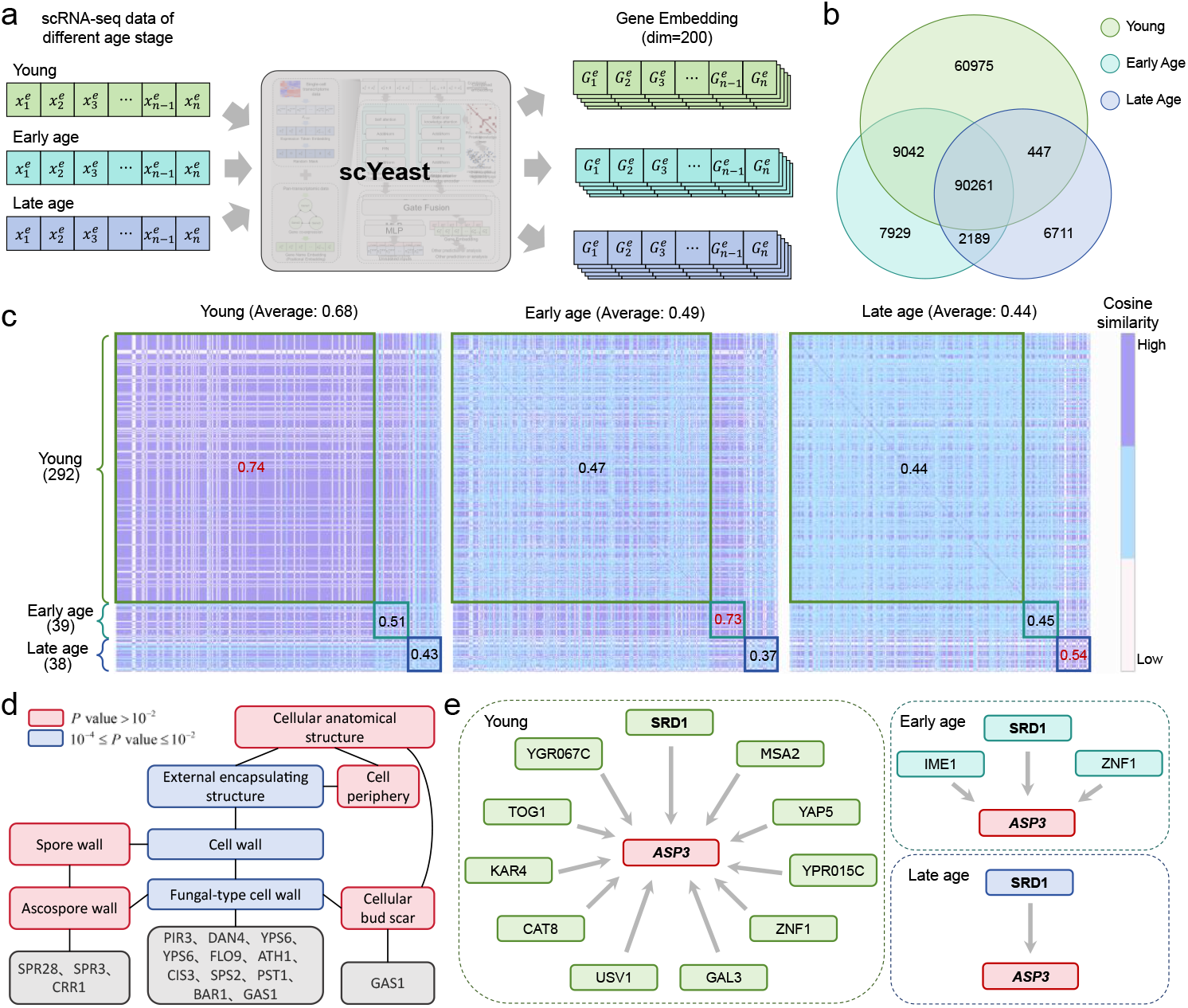
Inference of transcriptional regulatory relationships using scYeast. (a) Gene expression data from *Saccharomyces cerevisiae* at different age stages (young, early age, late age) are input into the scYeast model. The input data is embedded into 200-dimensional vectors, and the cosine similarity between these gene embeddings is calculated to infer regulatory relationships. (b) A comparative distribution plot of gene pairs with high similarity across the three age stages. (c) Heatmaps of the inferred important gene relationships at each age stage, clearly showing the changes and transitions in regulatory relationships with age. The mean similarity at each stage is calculated to reflect the changes in gene regulatory relationships at different ages. (d) GO enrichment analysis results for cellular components of important genes identified in the young stage. (e) A case study of *ASP3*, examining the changes in its inferred regulatory network with age, revealing a shift in the network’s focus as aging progresses.

First, we used scYeast to obtain gene embedding vectors for each sample. Gene pairs with a cosine similarity higher than 0.9 within each sample were defined as highly-correlated gene pairs, hinting a potentially strong interaction between them. Then, we extracted the highly-correlated pairs that were unique to each of the three age states. The results indicate that the number of highly-correlated gene pairs significantly decreased as cells age. A total of 160,725 highly-correlated pairs were extracted from the young state, whereas only 99,608 pairs remained in the late-aged state. Furthermore, the number of overlapping highly-correlated gene pairs between adjacent age stages was greater (Fig. 2b). Additionally, all genes appearing in the highly-correlated pairs were defined as “important genes.” We found 297 important genes exclusive to the young stage, while only 38 and 39 were exclusive to the early- and late-aged stages, respectively. This demonstrates that the intracellular transcriptional regulatory landscape in *S. cerevisiae* undergoes dynamic changes along with the cell states. To more intuitively display these age-dependent changes in gene regulation, we selected these important genes and plotted heatmaps for the three states (Fig. 2c), which clearly show a significant shift and remodeling of gene-gene regulatory relationships as cells age. To obtain more direct statistical features, we binned and calculated the average cosine similarity for the important genes exclusive to each age stage (Methods). It is evident that for each state, the average cosine similarity of its corresponding age-specific important genes is the highest. The overall average cosine similarity across all genes also decreases with aging (0.68 to 0.49 to 0.44), which reflects the changing trend of the co-expression relationships of genes within cells as age increases. GO enrichment analysis of the important genes from each stage revealed that among the genes exclusive to the young stage, 14 were significantly enriched (*P* value = 0.00715) in the “external encapsulating structure” under the cellular component annotation (Fig. 2d). This reflects the high dependency of young cells on cell wall functions, including growth, division, morphology maintenance, and stress response ^26^. As cells age, the function of the cell wall may gradually decline, leading to the disappearance of this significant enrichment. These findings fully demonstrate the rich information contained within scYeast’s gene embeddings and underscore its immense potential for inferring gene regulatory networks and their dynamic changes under various internal and external perturbations.

Further, we conducted a case study on *ASP3* (asparaginase II) to specifically analyze the inferred regulatory relationships. *ASP3* is a key enzyme in nitrogen metabolism and stress response, playing a crucial role in various biological processes ^27^. As cells age, the number of TFs that can significantly regulate *ASP3* decreases accordingly. From the perspective of expression levels, the correlation between the transcription factors and the expression levels of the target genes weakens over time (Fig. 2e). In young cells, the expression of *ASP3* is synergistically regulated by multiple transcription factors, ensuring a rapid response to environmental changes. However, as cells aging, the regulatory network tends to be simplified and relaxed, which may be due to the decreased transcription activity of TFs. This may hint that aging could remodel cellular functional states through a slow shift in the cell’s regulatory networks ^28^. While this regulatory simplification strategy might offer energy benefits, it could also reduce the adaptability of aging cells to environmental fluctuations and have potential negative impacts on other cellular functions ^29^.

### scYeast enables the prediction of various yeast phenotypes and identification of the related key genes

scYeast can also accurately predict various yeast phenotypes, i.e., the growth doubling time. To test if the prior knowledge-infused attention mechanism introduced in scYeast enhances phenotypic prediction ability, we trained an additional model. In this control model, the static, prior knowledge-based encoder was replaced with a standard transformer encoder, which was employed for phenotypic prediction in the same manner.

The network architecture for predicting yeast cell growth doubling time (regression) and different states (classification) is shown in Fig. 3a. New data is fed into the pre-trained model to generate gene embeddings, which then serve as the input for a downstream MLP network that performs the classification or regression task. Concurrently, to verify that the proposed pre-training network has effectively learned features from the unlabeled data, we also fed the raw data directly into several traditional machine learning algorithms for a comparative analysis of prediction results (Methods). The test results for the regression prediction of yeast growth doubling time are presented in Fig. 3b. It is evident that the prediction results from models incorporating the pre-trained embeddings outperform those from traditional machine learning algorithms, indicating that the pre-trained model has successfully learned useful information from large-scale unlabeled data, which in turn helps to improve the model accuracy in other predictive tasks. Furthermore, we observed that the pre-trained model incorporating prior knowledge surpassed the model without it, suggesting that the integration of prior knowledge during pre-training allows the model to further enhance its predictive accuracy. Under identical dataset partitioning (Methods), we further compared the effect of including prior knowledge in the fine-tuning network. The results show that the predictive performance of the model with added prior knowledge is superior overall to the one without, with a particularly significant advantage for samples with long doubling times.

**Figure 3.**
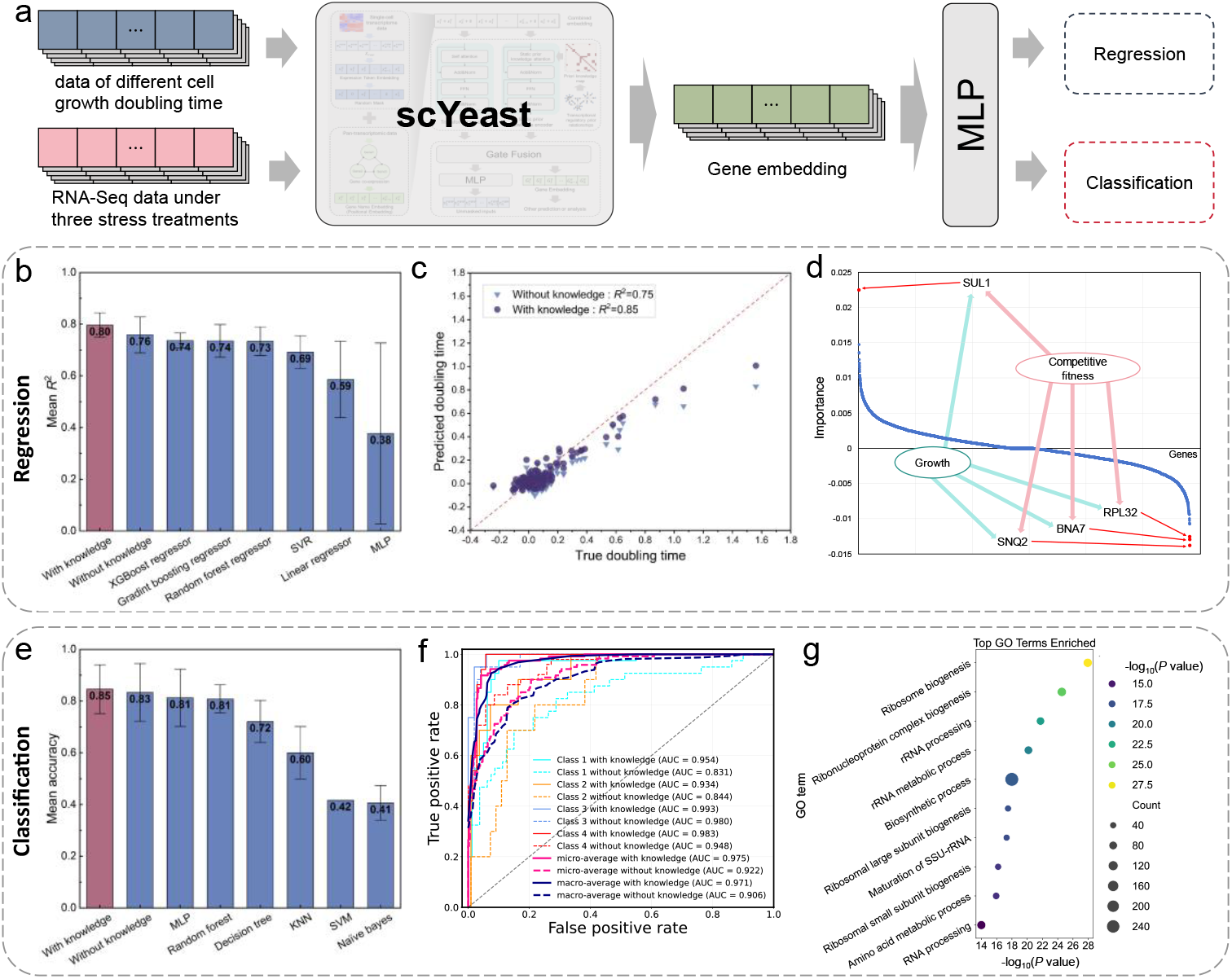
scYeast enables growth prediction and cell state classification tasks. (a) The network architecture for scYeast performing phenotype prediction tasks. The parameters of scYeast are fixed, and it is followed by an MLP for classification or regression prediction. (b) Comparison of results for the regression task of predicting growth doubling time. (c) A scatter plot comparing the prediction results on the test set for fine-tuned networks with and without the inclusion of prior knowledge, under the same dataset split. (d) The gradient distribution of key genes from the yeast growth prediction model, and the four key genes with the most significant gradients. (e) Comparison of results for the classification task of cell stress states, which are divided into four categories: isosmotic, hypoosmotic, glucose starvation, and amino acid starvation. (f) A comparison of the cell stress state classification results between the model with prior knowledge and the model without. The results are displayed using ROC curves, where solid and dashed lines distinguish the two types of models. The plot includes the individual ROC curves for each of the four classes and their average curve. (g) GO enrichment analysis results for the key genes identified by the cell stress state classification model.

Moreover, to derive biological insights, we interrogated this predictive model to identify key genes exerting the strongest influence on yeast growth doubling time. We calculated the gradient of the output (growth doubling time) with respect to each input variable. The results revealed that the absolute values of the gradients for four genes—SUL1, SNQ2, BNA7, and RPL32—were significantly larger than those of other genes (Fig. 3b). Therefore, we hypothesize that these four genes may be related to yeast growth doubling time. To assess the plausibility of this result, we examined annotations in the *Saccharomyces* Genome Database (SGD) ^30^, which links all four genes the keywords “competitive fitness” and “growth”, initially indicating that the regression model based on scYeast could identify genes pivotal to yeast growth phenotypes.

Similarly, the predictive results for the four-class classification task under different stress conditions are shown in Fig. 3e, with several traditional machine learning classification methods included as the reference. Consistent with those from Fig. 3b, the classification accuracy of the pre-trained network is higher than that of traditional algorithms, and the model incorporating prior knowledge outperforms the one without it. The ROC curve comparison in Fig. 3f between the pre-trained networks with and without prior knowledge further corroborates this conclusion. Subsequently, we calculated the gradients for the glucose starvation and amino acid starvation stress conditions with respect to each input variable within the four-class classification model to identify key gene sets for these two types of stress condition. GO enrichment analysis was performed on these gene sets, and the top ten enriched biological processes ranked by significance are shown in Fig. 3g. We found that they are significantly enriched in multiple ribosome-related processes (*P* value =1.17 ×10^−28^) and amino acid metabolic processes (*P* value =1.17 ×10^−16^). This is consistent with known biology, as yeast adaptation to glucose starvation involves specific changes in ribosomes, which cease to participate in protein synthesis. Likewise, under amino acid starvation stress, protein production is affected, leading to changes in ribosome-related processes ^31^.

### scYeast can predict temporal responses to gene perturbations

scYeast can serve as a foundational model to enable the prediction of responses to gene perturbations. Since scYeast has already integrated prior knowledge such as gene co-expression and transcriptional regulatory relationships during its pre-training, we adapted the GEARs ^32^ framework by using scYeast as the base model for predicting gene perturbation responses (Methods) (Fig. 4a). To predict the temporal responses under dynamic gene perturbation, the newly conceived network consists of three components: scYeast, a Gene Ontology (GO) graph network, and an MLP prediction network. First, the gene expression from the previous time point is used as input to scYeast to obtain a gene embedding representing the transcriptomic state at that moment. Subsequently, the perturbation information is fed into a graph convolutional neural network based on a GO relationship graph to obtain a high-dimensional embedding of the perturbation. Finally, the perturbation embedding and the transcriptomic state embedding are summed and input into an MLP for output prediction (Methods). Following this approach, we constructed ten independent perturbation networks, each corresponding to a unique gene (Table 1). Analysis of the training performance on the validation and test sets showed that each network reached its optimal validation performance within 15 epochs. Testing these ten optimal networks on the test dataset revealed that the *R*^2^ values ranged between 0.70 and 0.97 (Fig. 4b). This indicates that all networks were able to accurately capture the dynamic changes of these ten genes in response to the perturbations of transcription factor.

**Table 1.**
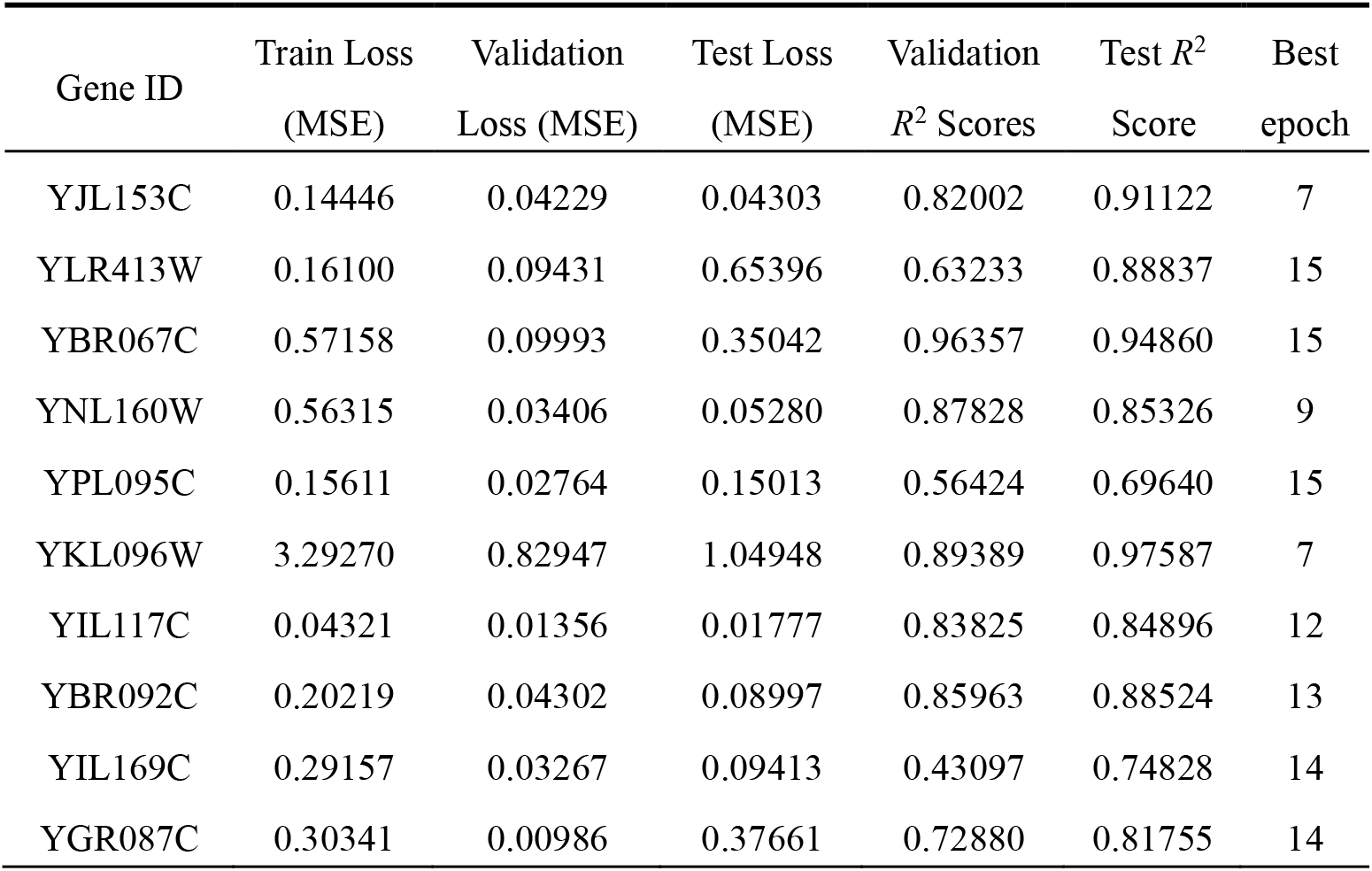
Training Information of Genes under the Perturbation of a Single TF.

**Figure 4.**
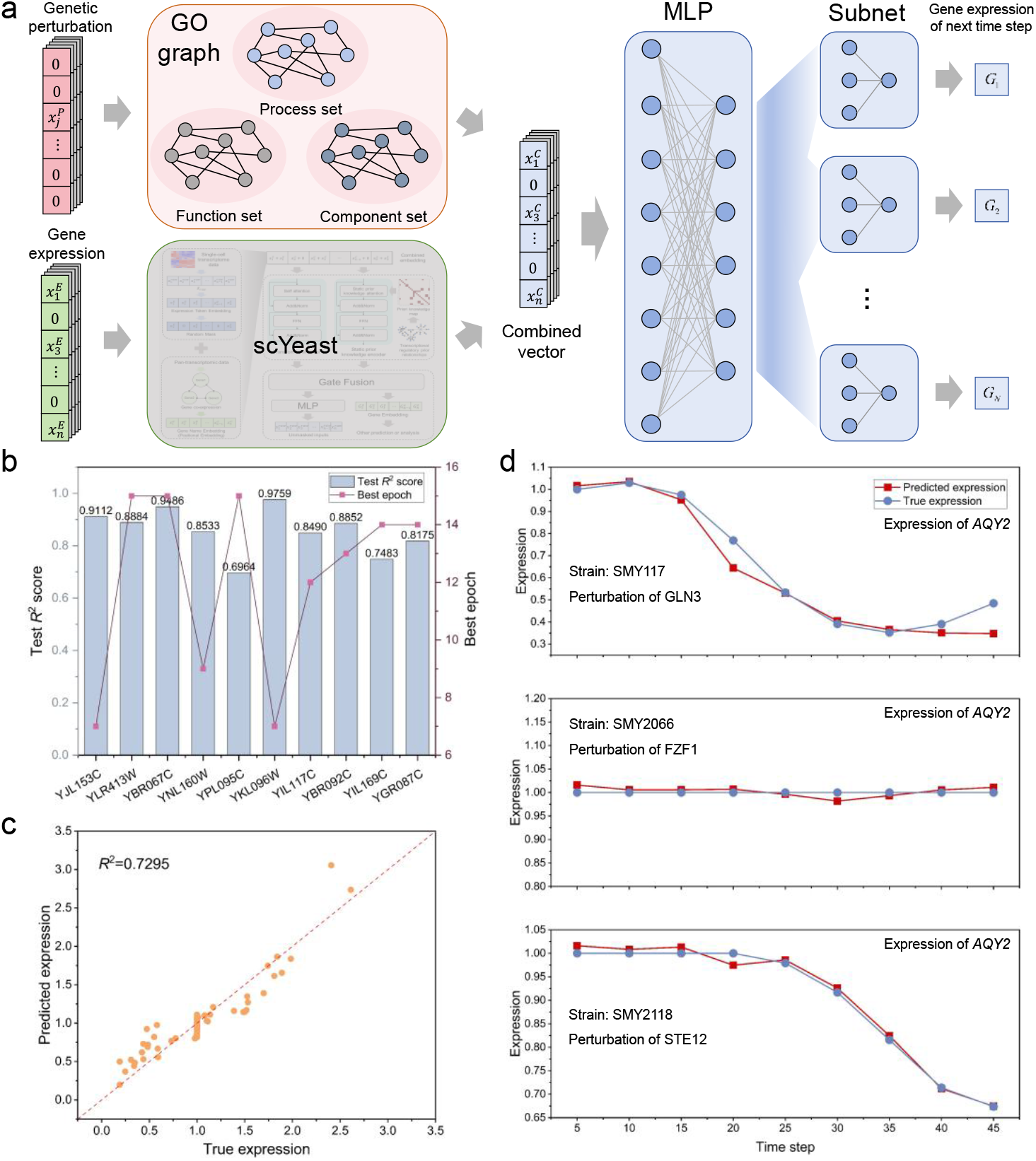
scYeast can predict the response of genes under the influence of single TF perturbations. (a) The input includes gene expression data at a specific time point and information about the transcription factor perturbation applied to the strain. These inputs are processed separately through the pre-trained scYeast model and a graph neural network to extract embeddings. The outputs from both are then concatenated and passed through multiple linear layers to predict the target gene expression level at the next time point under the given condition. (b) Training results for subnetworks targeting ten key genes. The bar chart represents the *R*^2^ scores of the subnetworks on the test set, while the curve indicates the training epoch at which the model achieved its optimal performance. (c) An *R*^2^ scatter plot on the training set for the subnetwork designed to predict the gene *AQY2*, after removing three types of perturbations from the training data. (d) A comparison of the predicted full time-series against the actual observed values under three perturbations (namely SMY117, SMY2066, and SMY2118) that were not included in the training. (SMY117, SMY2066, and SMY2118 represent different types of cellular perturbations, corresponding to three intervention experiments of TF—GLN3, FZF1, and STE12).

To further validate the model’s training performance, we selected *AQY2* as a case study and fine-tuned a dedicated prediction network for this gene. *AQY2* is an aquaporin ^33^ that enhances the permeability of the cell membrane to water. Studying *AQY2* helps to provide deeper insights into how yeast cells regulate water influx and efflux, and its role in osmoregulation and other physiological processes. In the public IDEA dataset, the temporal changes in gene expression following regulation by a transcription factor (TF) can be broadly categorized into three types: an S-shaped increase or decrease, a unimodal pulse, or remaining essentially stable (indicating the gene is not regulated by the stimulated TF) ^34^.

If the model truly captures the underlying regulatory relationships between TFs and target genes, it should be able to reproduce these typical dynamic profiles even when the time-series data for the perturbation of the corresponding TFs is absent. Based on this rationale, we entirely removed the time-series data from the intervention experiments of three TFs—GLN3, FZF1, and STE12 (strain numbers in dataset: SMY117, SMY2066, and SMY2118, respectively)—from the training data ^34^, and fine-tuned the network only on the remaining intervention conditions. This forces the model to rely on the learned regulatory priors and inter-genic temporal associations during the prediction phase, rather than performing simple regression. After 13 epochs of training, the optimal network achieved an *R*^2^ of 0.7295 on the held-out test set (Fig. 4c), indicating that the overall fit remained reliable. Subsequently, we reconstructed the complete temporal response of *AQY2* under the three aforementioned interventions, starting from time zero. The results are shown in Fig. 4d. Under GLN3 and STE12 interventions, the model-predicted expression of *AQY2* exhibited an inverse S-shaped curve, consistent with the experimental sequencing data ^34^. In contrast, under the FZF1 intervention, the predicted trajectory remained close to the baseline, accurately reproducing the stable “no significant response” pattern observed experimentally. This is consistent with biological priors obtained from the YEASTRACT database ^30^. In fact, GLN3 and STE12 are both confirmed to regulate *AQY2*, participating in the amino acid starvation response and the invasive growth signaling pathway in the cell cycle, respectively. In contrast, there is no known direct regulatory relationship between FZF1 and *AQY2*.

The above analysis demonstrates that scYeast could effectively mines the rich information embedded in transcriptomic data. The extracted transcriptional regulatory relationships could help to probe the dynamic correlations between TFs and their target genes in response to perturbations.

### scYeast can be transferred to proteomics data for growth prediction

As scYeast was mainly trained on single-cell transcriptomics, we further explored its potential applications on proteomic data through transfer learning (Fig. 5a). The proteomic data used was derived from a global proteome analysis of 4,699 *S. cerevisiae* single-gene knockout strains ^35^, in which 1,850 proteins were quantified using micro-flow liquid chromatography-tandem mass spectrometry (micro-flow LC-SWATH-MS), covering approximately 79% of the yeast coding genome. Due to the significantly lower gene dimensionality of proteomic data compared to transcriptomic data, we employed a zero-padding method to expand the proteomic data to the same dimension as the transcriptomic data, thereby fitting the input structure of the pre-trained model. Subsequently, the same random masking auto-regressive training strategy was applied to the proteomic data to generate protein-specific embeddings. Finally, global features of the embeddings were extracted via a pooling operation and input into an Extra Trees ensemble regression model to predict three types of model outputs collected from different literature sources: growth rate ^35^, ribosome occupancy of different genes (translation efficiency)^36^, and protein half-life ^37^ (Methods).

**Figure 5.**
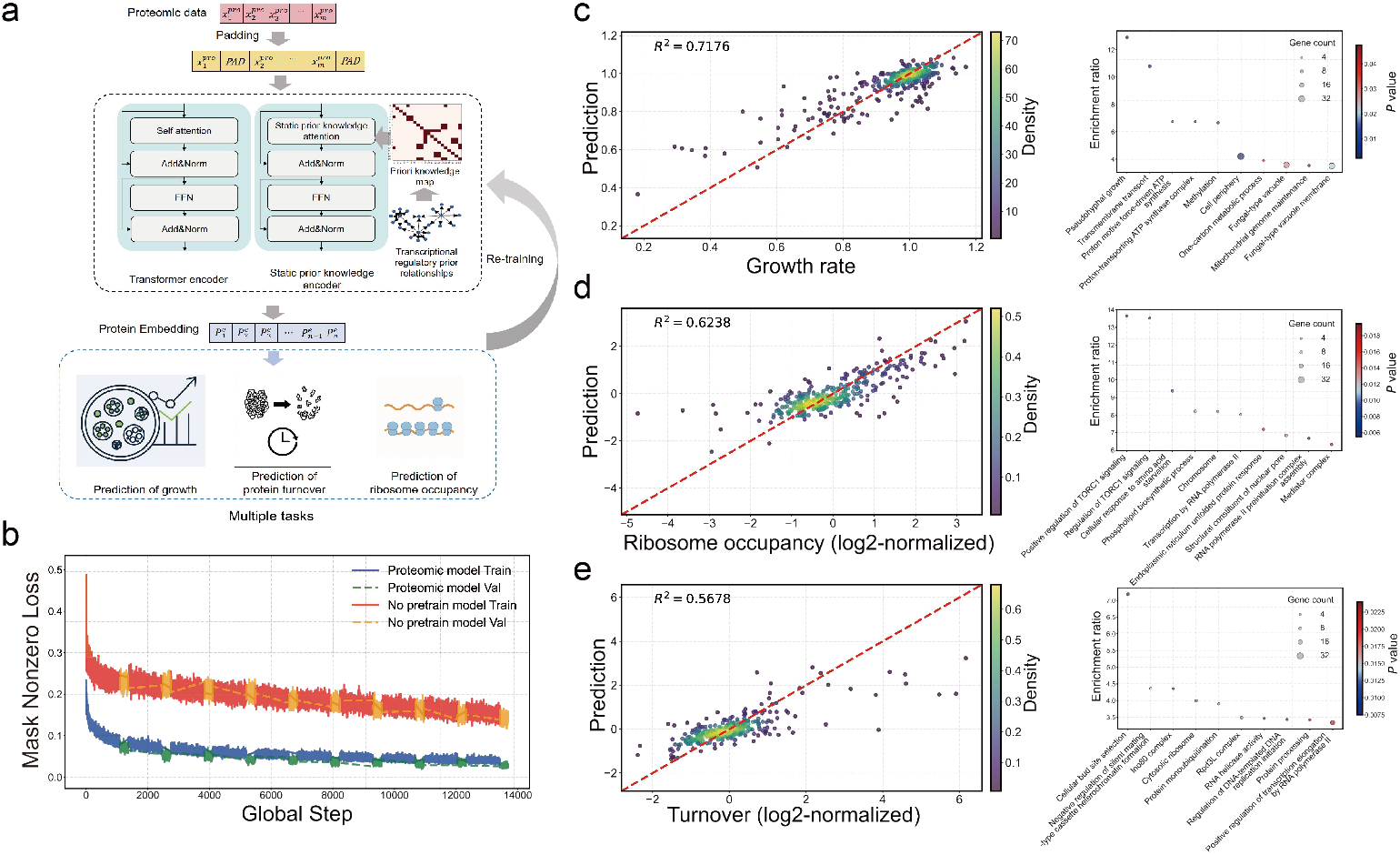
Transfer learning of scYeast to proteomics for phenotype prediction. (a) The network architecture for transfer learning of scYeast on proteomics data. (b) Iteration curves for transfer training with loaded scYeast transcriptome pre-trained parameters versus randomly initialized parameters. It is evident that the reconstruction error of the model with pre-trained parameters is lower than that of the model with randomly initialized parameters, demonstrating the effectiveness of the pre-trained model. (c) The *R*^*2*^ result for growth phenotype prediction using the proteomic scYeast model, and the corresponding Gene Ontology (GO) enrichment analysis of genes based on the model’s feature importance parameters. (d) The *R*^*2*^ result for ribosome occupancy prediction using the proteomic scYeast model, and the corresponding GO enrichment analysis of genes based on the model’s feature importance parameters. (e) The *R*^*2*^ result for protein half-life (turnover) prediction using the proteomic scYeast model, and the corresponding GO enrichment analysis of selected genes based on the model’s feature importance parameters.

To evaluate whether the original pre-trained model enhances the prediction performances of the newly generated model with proteomics as input, we compared the training processes of a model with loaded pre-trained parameters against one with randomly initialized parameters (Fig. 5b). The results indicated that the model with loaded pre-trained parameters had a significantly lower initial reconstruction loss at the masked sites compared to the control group, and the loss value after convergence was also significantly lower. This indicates that the knowledge learned by pre-trained model could effectively improve the training efficiency and prediction accuracy when replacing transcriptomics with proteomics as the input during transfer learning.

Finally, we used the pooled protein embedding features from the scYeast model as input for an Extra Trees regression model to predict phenotypic data (Fig. 5c-e). For all three predictions, the high R^2^ values could be achieved, demonstrating that protein embeddings from the newly fine-tuned scYeast effectively capture variations in protein abundance and enable precise, non-linear prediction of diverse phenotypes. Using the model’s feature importance scores, we selected top genes and performed Gene Ontology (GO) enrichment analysis to identify key biological pathways influencing growth, protein half-life, and gene translation, respectively. The GO enrichment analysis of proteins correlated to growth highlighted several pivotal processes, including “pseudohyphal growth” (*P* value = 0.006), a morphological adaptation mechanism of yeast under nutrient deprivation; “transmembrane transport” (*P* value = 0.009), essential for nutrient uptake and waste excretion; and “one-carbon metabolic process” (*P* value = 0.039), which supports nucleotide and amino acid synthesis, driving cell biomass accumulation and proliferation ^38^; GO enrichment analysis further identifies bioprocess related to ribosome occupancy, such as “positive regulation of TORC1 signaling” (*P* value = 0.005) and “cellular response to amino acid starvation” (*P* value = 0.006). These findings aligns with a recent study mentioning that TORC1, a key regulator of growth and metabolism, is modulated by amino acid availability, directly influencing ribosome activity, efficiency, and specificity ^39^. Lastly, for protein half-life, GO enrichment analysis highlighted processes such as “protein monoubiquitination” (*P* value = 0.016) and “protein processing” (*P* value = 0.023), consistent with the role of the ubiquitin-proteasome system as a critical pathway for protein degradation in eukaryotes, affecting protein stability and turnover ^40^. Collectively, through transfer learning, the model effectively captures rich representations from proteomic data, yielding complementary mechanistic insights into correlations between protein abundance and various phenotypes.

## Discussion

In this study, we have introduced scYeast—the first foundational single-cell model for yeast that deeply integrates prior regulatory knowledge. scYeast leverages an asymmetric parallel architecture to embed prior transcriptional regulatory information deeply into the Transformer’s attention mechanism, yielding gene embeddings for each sample that fully consider both prior knowledge and data-driven features. Based on these gene embeddings, it is possible to infer condition-specific transcriptional regulatory relationships and extract biological insights. Ablation studies demonstrated that the pre-trained model, which integrates regulatory prior knowledge, can effectively learn feature information from large-scale, unlabeled single-cell transcriptomics data. This, in turn, enhances the accuracy of cellular phenotype predictions, such as doubling time and state classification, while also enabling the analysis and identification of key gene sets that critically influence these phenotypes. The potential of scYeast as a base model was also validated, as it exhibited outstanding performance in predicting gene perturbation responses, accurately simulating the dynamic changes of genes under different perturbations. Finally, by applying transfer learning with scYeast to proteomics data, we further confirmed the effectiveness of the pre-trained model, successfully applying it to proteomics-based growth prediction and key gene discovery.

However, the prior knowledge in scYeast is based on a static relationship matrix derived from regulatory databases. This means that regardless of the conditions under which the input transcriptomics data were obtained, the transcriptional regulatory prior knowledge within the model remains fixed. In reality, studies have shown that yeast transcriptional regulatory relationships are dynamic ^41,42^, varying with different growth conditions and developmental stages. Therefore, allowing the transcriptional regulatory prior knowledge to adjust dynamically with changes in the input data could further improve the model’s accuracy. In the future, integrating more diverse forms of prior knowledge into the model could also be considered. At the same time, the scale of single-cell transcriptomics data for yeast has not yet reached its full potential. As data volume continues to accumulate, it will be feasible to construct pre-trained models with a larger number of parameters, thereby enhancing the precision of simulating transcriptional regulatory relationships, extracting more feature information, and further improving model accuracy. Furthermore, while this work has highlighted several typical application scenarios, as a foundational pre-trained model, scYeast can, in fact, be adapted for many other computational tasks, which can be further explored by wider yeast community.

In summary, scYeast, as the first large-scale pre-trained model for yeast that deeply integrates prior biological knowledge, has demonstrated strong generalizability and practical utility. This foundational model not only offers a powerful, reliable tool for dissecting transcriptional regulation in yeast but also establishes a framework for seamlessly integrating diverse biological datasets, thus significantly accelerating our mechanistic understanding of yeast cellular metabolism.

## Methods

### Dataset preparation

The datasets required for the construction and training of scYeast are described below. Each dataset underwent quality control and preprocessing.

1. **Transcriptional regulatory prior knowledge** The prior knowledge of transcriptional regulation required by scYeast was obtained from the YEASTRACT database ^30^. The “documented targets” for each transcription factor were selected, which represent regulatory relationships supported by literature evidence. This approach ensures a high-quality, evidence-based prior knowledge matrix of regulatory interactions, comprising approximately 170,000 regulatory relationships in total.
2. **Pan-transcriptome data** The *Saccharomyces cerevisiae* pan-transcriptome data was composed of 969 high-quality transcriptomes, covering 4,977 core genes and 1,468 accessory genes ^24^. The Spearman correlation coefficient for each gene across different strains was calculated based on this dataset, serving as the basis for inferring co-expression relationships in *S. cerevisiae*.
3. **Single-cell transcriptomics data** The single-cell transcriptomics data refers to the data required for the pre-training of scYeast were primarily composed of two datasets: GSE125162 ^43^ and GSE242556 ^44^. In GSE125162, data were compiled from wild-type controls and 11 different genotypes under various environmental conditions, and expression states were determined by sequencing 38,285 single cells. In GSE242556, continuous sampling of *S. cerevisiae* without metabolic labeling was performed, and the expression of 173,361 individual cells was measured. Quality control was performed on the obtained single-cell datasets, and cells with total read counts below 200 were removed. Ultimately, a dataset of approximately 210,000 cells was constituted for the pre-training of scYeast.
4. **Yeast aging transcriptomics data** Transcriptomics data of single yeast cell at different age conditions were sourced from GSE210032 ^25^. This dataset includes 37, 43, and 45 single cells from the 2-hour (young), 16-hour (early-aged), and 36-hour (late-aged) age groups, respectively. On average, 2,202 genes were detected per cell, accounting for approximately one-third of the total genes in *S. cerevisiae*. To ensure the completeness of the analysis by scYeast, missing gene expression values were imputed with zeros.
5. **Transcriptomics data for growth prediction** To predict yeast cell growth phenotypes from transcriptomics data, a dataset comprising transcriptomes of 1,143 single-gene knockout strains was collected ^45,46^. The dataset comprises the transcriptomic data for each gene knockout together with the corresponding cell doubling times under specific conditions. The cell doubling time was used as the prediction label to simulate and predict yeast growth phenotypes from transcriptomic data.
6. **Stress condition data** The data for classifying different yeast stress states were obtained from GSE201386 ^47^. Using a full-length-based single-cell RNA sequencing method, 117 yeast cells were sequenced under four stress conditions (isosmotic, hypoosmotic, glucose starvation, and amino acid starvation). This data was used to perform classification prediction of these four cellular stress states.
7. **Time-series perturbation data** The yeast time-series perturbation dataset was sourced from IDEA ^34^. This dataset was constructed by independently inducing hundreds of transcription factors (TFs) and measuring the time course of the resulting gene expression responses in budding yeast. Over 200 TF induction experiments are encompassed in IDEA, each followed by rapid changes in genes directly regulated by these TFs and subsequent changes in indirectly regulated genes. Time-series data on the differential expression of the complete transcriptome of *S. cerevisiae* following perturbation by a single TF were provided by the IDEA dataset, typically spanning eight time points. However, due to the non-uniform distribution of time points in the IDEA dataset, a radial basis function interpolation method with a Gaussian kernel was employed to interpolate the data within a 45-minute window at 5-minute intervals.
8. **GO relationship matrix** For the time-series perturbation prediction, the perturbation relationship graph was derived from the Gene Ontology (GO) database ^48,49^. GO comprises three domains: Biological Process, Molecular Function, and Cellular Component. All GO IDs for each gene across the three domains were retrieved, and connections were created between genes sharing the same GO IDs. The more GO IDs shared, the greater the connection weight. This process was used to construct an adjacency matrix of inter-genic GO relationships, which served as the perturbation relationship graph for the time-series perturbation prediction
9. **Proteomics data for growth prediction** Proteomics and phenotype data for prediction The proteomics data were adopted from a global proteome analysis of 4,699 *Saccharomyces cerevisiae* non-essential gene knockout strains, which quantified 1,850 proteins ^35^. The growth rate data for 4,550 of these strains were also adopted from the same study ^35^. Data for the other prediction tasks were sourced separately: ribosome occupancy data were derived from McManus et al. ^36^, and protein half-life data were from Martin-Perez and Villén ^37^.

### Static prior knowledge attention

In scYeast, to more deeply integrate prior knowledge within the Transformer framework, a static graph attention method was employed. First, the transcriptional regulatory prior knowledge is stored in the form of an adjacency matrix *P*. Thus, *P* is a square matrix where the number of rows and columns equals the number of genes, and a value of 1 in *P* indicates that a regulatory relationship exists between the genes corresponding to the row and column. Subsequently, considering the attention calculation formula in the Transformer framework:

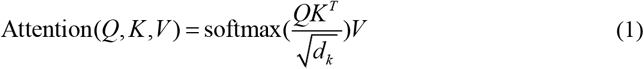

where Q, K, and V are the query, key, and value matrices, respectively. The term can be defined as the attention weight matrix, which is also a square matrix with dimensions equal to the number of variables. When multiplied by the value matrix *V*, it represents the attention relationships between variables. Therefore, the attention weight matrix was replaced with the prior knowledge adjacency matrix *P*, signifying that the attention relationships between variables can be guided by the provided prior knowledge.

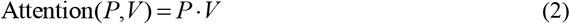

Here, *P* represents static prior knowledge; thus, the parameters within the *P* matrix are not updated during the model training process.

### Gene name embedding

In scYeast, the gene name embedding is determined by gene co-expression relationships. First, a Spearman correlation analysis was performed on the expression levels of every gene pair using the pan-transcriptome data. These correlations were then filtered using a threshold of 0.5. That is, when the Spearman correlation coefficient is greater than 0.5, the two genes are considered to have a significant co-expression relationship, and were saved as a gene pair. Subsequently, each gene pair is treated as a “sentence” in natural language processing. If a gene has no identified co-expression relationships, it forms a sentence consisting of a single “word.” The Word2Vec ^50^ method was then used to obtain a 200-dimensional embedding vector for each gene. In scYeast, this gene name embedding replaces the conventional positional embedding, and the position of each gene in the input data must be kept constant.

### Expression token embedding

For the gene expression value embedding module, scYeast references the continuous embedding strategy from scFoundation ^5^ and adapts it to the specific characteristics of the yeast dataset. For the input single-cell transcriptomics data, the raw expression value x was first mapped to a high-dimensional vector representation. Specifically, non-zero expression values generate dynamic attention weights through a two-stage linear transformation. The first MLP layer projects the scalar value into a hidden space, and after LeakyReLU activation, a second MLP layer further learns non-linear associations across dimensions. The formulas are as follows:

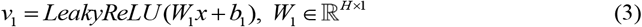

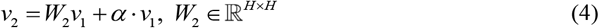

where *α* (initialized to 1.0) is a learnable scaling factor. The scores are then obtained through Softmax normalization:

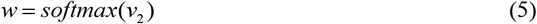

Finally, a continuous embedding vector is generated by a weighted summation based on a predefined embedding table:

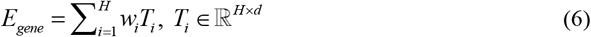

where *H* is the predefined number of bins (default is 10) and is the preset embedding vector dimension. For special tokens (zero values, masked positions, padding symbols), the model initializes separate embedding layers, *E*_zero_, *E*_mask_ and *E*_pad_, respectively. In the implementation, the positions of these special tokens are located by index and replaced with their corresponding embedding vectors. This continuous embedding strategy avoids the information loss associated with traditional binning of continuous data at fixed intervals and eliminates the problem of selecting an appropriate bin size. The gene-specific embedding and the expression value embedding are fused via element-wise addition to form the final input representation.

### Pre-training of scYeast

scYeast adopts a pre-training strategy typical of large models. During the pre-training phase, scYeast employs a self-supervised, auto-regressive training process following random masking. Specifically, when single-cell transcriptomics data were received as input by the model, the gene name embeddings are not masked. Only 15% of the expression token embeddings are randomly masked. The model’s output is the original single-cell transcriptomics data itself, thereby achieving auto-regressive training.

During pre-training, scYeast utilizes the Huber loss function. Considering the high sparsity of single-cell transcriptomics data, a large number of zero values could potentially impact the loss function and the training process during auto-regressive reconstruction. Furthermore, to emphasize the reconstruction performance on the masked portions, scYeast employs a composite loss function, as shown in the formula:

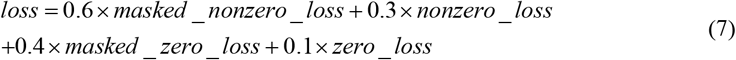

The training parameters for the model were set as follows: a batch_size of 16, the AdamW optimizer with a weight decay of 0.01, a learning rate set to 5e-5, and a maximum of 10 epochs. The model was constructed and trained within the PyTorch environment, utilizing GPU parallel computing for acceleration.

### Predicting gene perturbation response

With scYeast serving as a base model, we predict the gene expression at the next time point by using the gene expression data from the previous time point as input, thus achieving single-step, time-series prediction of gene expression response. To mitigate the impact of the numerous zero values in the IDEA dataset, an exponential transformation was applied to the data. The input data consists of two parts. First, the gene expression data at a specific time point following TF perturbation is fed into scYeast to obtain context-aware gene embeddings. Second, the perturbation vector is input into the composite GO relationship graph to generate a perturbation embedding. These two embeddings are then summed and passed through multiple linear layers for progressive dimensionality reduction to predict the expression of a specific gene at the next time point.

### Transfer learning for proteomic data

The knowledge learned by scYeast during its transcriptomic pre-training was transferred to the pre-training of a model for proteomics data. Since the dimensionality of proteomics data is much smaller than that of transcriptomics data, the dimensionality of the proteomics data was expanded to match that of the transcriptome by padding the gene positions that had no protein match. This ensured compatibility with the model’s architecture. The new model adopts the same architecture as the transcriptomic scYeast. During its pre-training, two schemes were employed to validate the effectiveness of transferring knowledge from the transcriptomic network to the proteomic network: one involved loading the parameters from the transcriptomic model, while the other used randomly initialized parameters. This model also utilized a self-supervised, auto-regressive training process following random masking. In the model’s forward propagation architecture, the weights of the padded positions in the attention matrix were masked out to prevent the token information from non-protein-matched sites from interfering with the training.

### Prediction of growth rate, ribosome occupancy, and protein half-life based on proteomics data

The embedding vectors corresponding to the proteomic data, obtained from the pre-trained model, were processed through max-pooling and min-pooling, then concatenated to form feature vectors. These vectors were input into an Extra Trees (n=200) model to predict the growth rate of strains in YPD medium^35^, as well as experimentally measured ribosome occupancy ^36^ and protein half-life^37^. The training was conducted over multiple rounds using a k-fold (k=10) training scheme. For comparison, the raw proteomic abundance information of the strains was also input as feature vectors into the Extra Trees model for prediction using the same scheme. The growth rate prediction was mainly used to analyze the effect of proteins on growth rate. The ribosome occupancy and protein half-life data were used as response variables to analyze the impact of gene knockouts on translation efficiency and protein turnover rate.

Key genes or gene knockouts were identified by screening based on the intrinsic feature parameters of the Extra Trees model. During the prediction process, the Extra Trees (n=200) model, trained on the pooled features of the proteomic embeddings, automatically assesses the predictive contribution of each input feature to the target variable according to the built-in Gini importance calculation method of the scikit-learn framework. This importance score quantifies the extent to which a feature affects the model’s output. These features were ranked by their importance scores, and the top 100 genes or gene knockouts were selected for subsequent GO enrichment analysis using goatools ^49^.

## Acknowledgement

This work was financially supported by the National key research and development program of China (2020YFA0908300), Shanghai Municipal Science and Technology Major Project, and grant 22208211 and 22378263 from the National Natural Science Foundation of China (NSFC). Portions of this work were used in the PhD thesis of the first co-author. This manuscript benefited from language polishing using AI assistant; all scientific content (study design, data, analyses, and conclusions) was conceived, verified, and is fully the responsibility of the authors.

## Code and data availability

The scYeast is available at the repository: https://github.com/hongzhonglu/scYeast. Data is available at https://figshare.com/articles/dataset/data_for_scYeast/30109528.

## Author contributions

H.L. and X.Y. conceived the study, supervised the work, provided guidance, and obtained the funding for this study. X.F. proposed the modeling framework, and conducted model pretraining and debugging. W.L. implemented and debugged the first three applications of the pretrained model, analyzed experimental results, and prepared the figures. L.X. implemented and debugged the fourth application of the model. All authors contributed to manuscript revision.

